# The thiol methyltransferase activity of TMT1A (METTL7A) is conserved across species

**DOI:** 10.1101/2023.11.17.567538

**Authors:** José M. González Dalmasy, Christina M. Fitzsimmons, William J.E. Frye, Andrew J. Perciaccante, Connor P. Jewell, Lisa M. Jenkins, Pedro J. Batista, Robert W. Robey, Michael M. Gottesman

## Abstract

Although few resistance mechanisms for histone deacetylase inhibitors (HDACis) have been described, we recently demonstrated that TMT1A (formerly METTL7A) and TMT1B (formerly METTL7B) can mediate resistance to HDACis with a thiol as the zinc-binding group by methylating and inactivating the drug. TMT1A and TMT1B are poorly characterized, and their normal physiological role has yet to be determined. As animal model systems are often used to determine the physiological function of proteins, we investigated whether the ability of these methyltransferases to methylate thiol-based HDACis is conserved across different species. We found that TMT1A was conserved across rats, mice, chickens, and zebrafish, displaying 85.7%, 84.8%, 60.7% and 51.0% amino acid sequence identity, respectively, with human TMT1A. Because TMT1B was not found in the chicken or zebrafish, we focused our studies on the TMT1A homologs. HEK-293 cells were transfected to express mouse, rat, chicken, or zebrafish homologs of TMT1A and all conferred resistance to the thiol-based HDACIs NCH-51, KD-5170 and romidepsin compared to empty vector-transfected cells. Additionally, all homologs blunted the downstream effects of HDACi treatment such as increased p21 expression, increased acetylated histone H3, and cell cycle arrest. Increased levels of dimethylated romidepsin were also found in the culture medium of cells transfected to express any of the TMT1A homologs after a 24 h incubation with romidepsin compared to empty-vector transfected cells. Our results indicate that the ability of TMT1A to methylate molecules is conserved across species. Animal models may therefore be useful in elucidating the role of these enzymes in humans.

## 1. INTRODUCTION

Methyltransferases (MTases) are a family of enzymes that transfer methyl groups to substrates. In most methyltransferases, S-adenosyl-L-methionine (SAM) acts as the methyl group donor, with S-adenosylhomocysteine (SAH) as a metabolic byproduct [1]. Substrates for methyltransferases are diverse and include small molecules, proteins, lipids, and nucleic acids [2]. MTases can be classified according to the molecule accepting the methyl group, with oxygen (*O*-), nitrogen (*N)*- or carbon (*C*-) MTases being most common, although a few thiol (*S*-) MTases are known to exist and some MTases can methylate more than one type of acceptor molecule [3]. Of the known human MTases, only a few are known to metabolize drugs and these include catecholamine O-methyltransferase (COMT), which can metabolize levodopa and thiopurine S-methyltransferase (TPMT), which metabolizes thiopurines used in the treatment of leukemia [4].

Another MTase linked to drug methylation is thiol methyltransferase (TMT), the existence of which was hypothesized over 60 years ago when it was found that incubating liver microsomes from humans and animals with compounds such as dimercaprol, 2-mercaptoethanol, and hydrogen sulfide led to their methylation [5]. TMT activity was also reported in the membranes of erythrocytes and was responsible for the methylation of the drugs captopril and 7-alpha thiospironolactone [6]. Recently, it was determined that METTL7A and METTL7B are the human TMTs that play this role, as both can methylate captopril and thiospironolactone, as well as other thiol-containing molecules including penicillamine and dithiothreitol [7, 8]. METTL7A and METTL7B were renamed TMT1A and TMT1B, respectively, to reflect their activity. We discovered that TMT1A was overexpressed in a series of cell lines selected for resistance to romidepsin, a histone deacetylase (HDAC) inhibitor (HDACi) [9]. Romidepsin is a prodrug that harbors a disulfide bond that must be reduced to yield the active form of the molecule. Activated romidepsin contains a thiol group that coordinates with a zinc ion to reversibly inhibit class I HDACs and methylation of the thiol group inactivates the drug [10]. We found that TMT1A overexpression confers resistance to HDACis with a thiol as the zinc-binding group, while TMT1B has a narrower substrate range [9]. We showed that TMT1A was able to methylate and inactivate the zinc-binding thiol of activated romidepsin, thus leading to drug resistance [9].

Animal model systems are frequently used to investigate the physiological function of proteins. While TMT1B is not conserved, TMT1A is found in many different model species, including the zebrafish, chicken, rat, and mouse. To determine if animal models could be used to gain insight into the role of TMT1A and TMT1B, we first needed to know if their functions are conserved across various species. By overexpressing TMT1A variants in HEK293 cells, we found that TMT1A proteins from different species were all able to confer resistance to thiol-based HDACis to varying degrees, suggesting that this function of TMT1A is conserved and that animal model systems could indeed be used to elucidate the physiological role of these methyltransferases.

## 2. MATERIALS AND METHODS

### 2.1. Chemicals

Romidepsin was obtained from Selleck Chem (Houston, TX). Belinostat was purchased from ChemieTek (Indianapolis, IN). KD 5170 and NCH-51 were from Cayman Chemical (Ann Arbor, MI). Dichloro-α-methylbenzylamine (DCMB, LY78335) was from Santa Cruz Biotechnology (Dallas, TX).

### 2.2. Cells

HEK-293 cells (ATCC, Manassas, VA) were maintained in Eagle’s minimum essential medium supplemented with 10% FBS, 2 mM glutamine and Pen/Strep. Cells were transfected with empty pcDNA3.1 plasmid or this plasmid containing human *TMT1A* (*Homo sapiens (Hs)*, NM_014033.4), mouse *Tmt1a1* (*Mus musculus (Mm)*, NM_027334.3), rat *Tmt1a* (*Rattus norvegicus (Rn)*, NM_001037355.1), zebrafish *tmt1a.1* (*Danio rerio (Dr)*, NM_001080039.1) or chicken *TMT1A* (*Gallus gallus (Gg)*, XM_424479.6), all with a C-terminus FLAG tag epitope (all from Genscript, Piscataway, NJ). Transfections were performed using Lipofectamine 2000 (ThermoFisher Scientific, Waltham, MA) according to the manufacturer’s instructions. Single cell clones were selected and maintained in 1 mg/mL G418 antibiotic (Mediatech/Corning, Manassas, VA).

### 2.3. Cytotoxicity assays

Cells were plated (5,000 cells per well) in opaque, white-walled 96-well plates and were allowed to attach overnight before treating with drug. The following day, drugs were added to plates in triplicate and incubated for 3 days before being developed by adding CellTiterGlo (Promega, Madison, WI) per manufacturer’s instructions. Plates were read on a Tecan Infinite M200 Pro (Tecan Systems, Inc., San José, CA). The drug concentration that inhibited 50% of cell growth (GI_50_) was determined from concentration curves.

### 2.4. Immunoblot analysis

Whole cell lysates were extracted from cell pellets with RIPA buffer (50 mM Tris – HCl [pH 7.4], 1% Triton X-100, 10% glycerol, 0.1% SDS, 2 mM EDTA, and 0.5% deoxycholate, 50 mM NaCl) containing protease inhibitor cocktail (cat# 5871, Cell Signaling Technology, Danvers, MA). To inhibit residual histone deacetylase (HDAC) activity after lysing cells, 500 nM trichostatin A was added to the RIPA buffer when examining histone acetylation. Following sonication, samples were centrifuged at 9500 x g for 10 minutes and soluble proteins in the supernatant were collected. Protein concentrations were determined using Bio-Rad Protein Assay Dye Reagent (Bio-Rad, Hercules, CA) according to the manufacturer’s instructions. Approximately 30 μg of protein lysates were separated by SDS-PAGE and transferred to nitrocellulose membranes. The membranes were probed with antibodies to detect FLAG tag (Millipore-Sigma, Burlington, MA cat# F1804-200UG), histone H3 (Millipore-Sigma, cat# 05-499), pan-acetyl histone H3 (cat# 06-599, Millipore-Sigma), beta-actin, (generated in mouse cat# 3700, or rabbit cat# 4970, Cell Signaling) and p21 (cat# 2946, Cell Signaling). After incubation with primary antibodies, membranes were incubated with fluorescently tagged IRDye secondary antibodies (LI-COR, Lincoln, NE, USA), and bands were visualized using the Odyssey CLx imaging system (LI-COR).

### 2.5. Cell cycle analysis

Cells were plated and allowed to attach overnight before being treated with the desired concentration of HDACi for 24 hours. Cells were then trypsinized, centrifuged, and the supernatant was removed. Equal volumes of staining solution (0.05 mg/ml propidium iodide with 0.1% Triton-X) and RNAse A solution (200 U/mL RNAse A in deionized water) were added to cells and incubated for 20 minutes. Samples were read on a BD FACSCanto Flow Cytometer (BD Biosciences) with propidium iodide fluorescence detected with excitation at 488 nM and a 585 nm bandpass filter. A minimum of 10,000 events per sample were collected. Modfit LT for Mac version 5.0 (Verity Software House, Topsham, ME) was then used to determine the percentage of cells in each phase of the cell cycle. Statistically significant differences between phases of the cell cycle were determined by a one-way ANOVA test with a post hoc Dunnett test for multiple comparisons.

### 2.6. LC-MS to detect methylated romidepsin

Plated cells were incubated with 10 μM romidepsin for 24 h after which the culture medium was centrifuged to remove any cellular debris. A 10 μL volume was injected for each sample on an Exion liquid chromatography system (SCIEX, Framingham, MA) coupled to an X500B QTOF mass spectrometer (SCIEX, Framingham, MA). Samples were separated on a 2.1 x 50 mm Aeris 3.6 μm WIDEPORE XB-C8 200 Å column (Phenomenex, Torrance, CA) at 0.3 mL/min using a 4 min linear gradient from 98% A (0.1% formic acid) to 40% B (acetonitrile with 0.1% formic acid) followed by a high organic wash and low organic column re-equilibration. Data were acquired in positive ionization mode across the mass range of 100-800 m/z, while data dependent acquisition MS/MS was performed across the mass range of 50-750 m/z. SCIEX OS software (SCIEX, Framingham, MA) was used for quantitation of extracted ion chromatogram peak areas.

### 2.7. Predicted protein modeling and protein alignment

Protein amino acid sequences (accession codes listed above) were obtained from the National Center for Biotechnology Information (NCBI) and predicted protein structures were obtained using AlphaFold v2.3.2 CoLab notebook [11]. Predicted three-dimensional structures were downloaded and visualized in Pymol (The PyMOL Molecular Graphics System, Version 2.0 Schrödinger, LLC.). Two-dimensional sequence alignment and phylogenetic tree analysis was constructed in Jalview [12] with Clustal Omega [13, 14] using the default parameters.

## 3. RESULTS

### 3.1. Sequence comparison of TMT1A homologs

The *TMT1A* gene is conserved across all the species we examined. While humans, rats and chickens have one copy of this gene, mice have three copies, *Tmt1a1, Tmt1a2* and *Tmt1a3*. Zebrafish also have three genes that are tandem duplications, *tmt1a.1, tmt1a.2* and *tmt1a.3*. Since the TMT1B gene is not conserved— chickens and zebrafish lack the gene—we chose to focus on conserved homologs of TMT1A to determine if its function was retained across species.

Amino acid alignment of the different TMT1A homologs showed high conservation among species (Figure 1A). In particular, we show that the GxGxG domain, a hallmark binding domain for SAM-dependent methyltransferases [15] (noted by red asterisks and the red letters at positions 78, 80 and 82 in human TMT1A) is conserved. Additionally, the aspartic acid at position 98 (denoted by a green asterisk) also predicted to bind SAM [7, 16], is conserved as well. Furthermore, we find that there are high conservation values and consensus at each amino acid. An examination of the amino acid sequence identity among the proteins (Fig. 1B) revealed that mouse and rat TMT1A have the closest sequence identity with humans at 85.7% and 84.8%, respectively, while chicken TMT1A has 60.7% identity and zebrafish only 51% identity. In addition, a phylogeny of TMT1A homologs indicates that human, mouse, and rat form one cluster, suggesting a conservation of methylation activity, while chicken and zebrafish are more divergent (Figure 1C). Three-dimensional modeling of human TMT1A (Fig. 1D) as well as mouse, rat, chicken, and zebrafish TMT1A (Fig. 1E) shows that all of these homologous proteins are predicted to adopt the conserved seven-beta-strand (7BS) MTase fold and show good structural alignment of the conserved *S*-adenosyl methionine (SAM)-binding domain and residues.

**Fig. 1.**
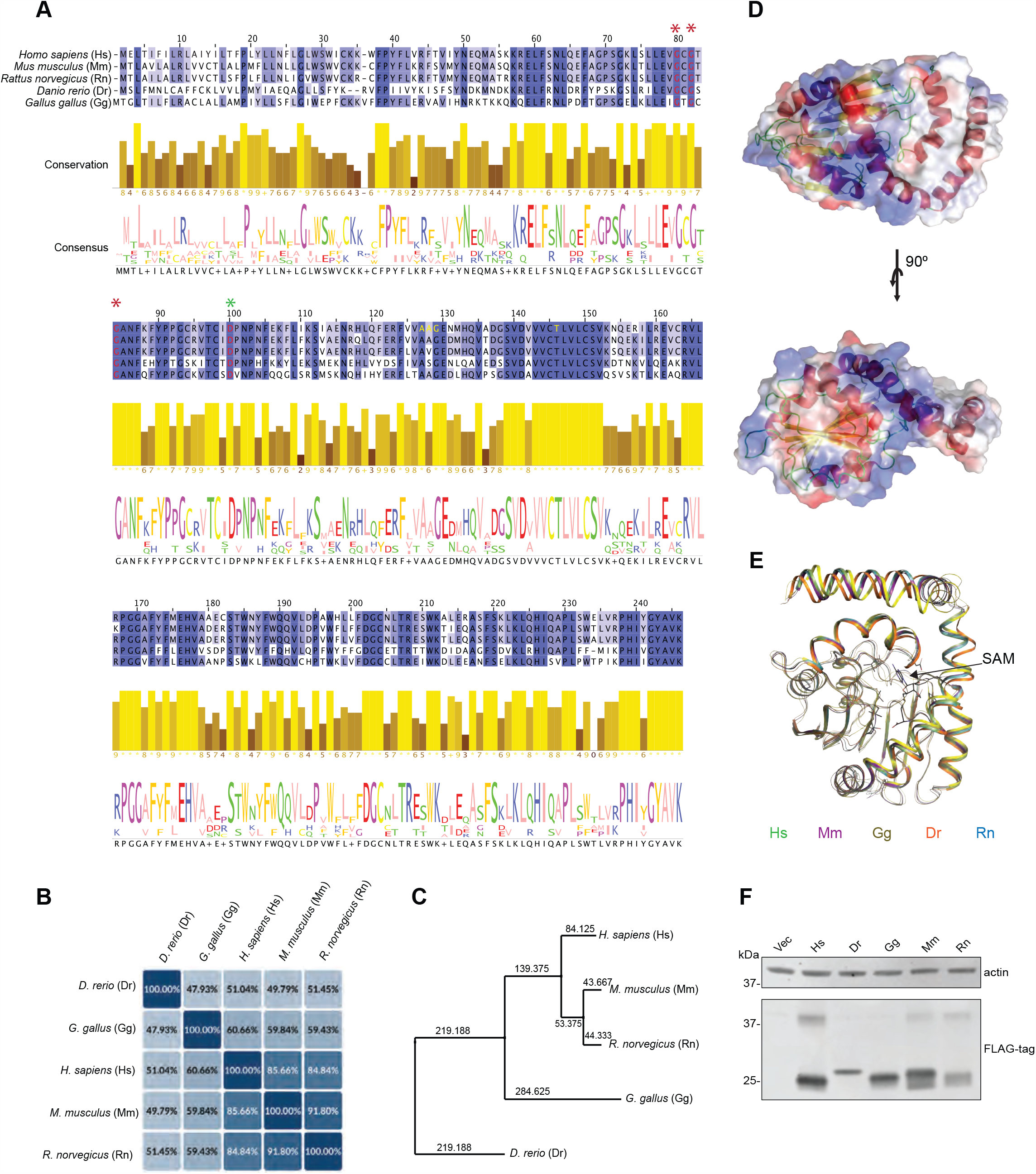
Comparison of amino acid sequence of TMT1A isoforms (A) Amino acid sequence alignment of human (Hs), mouse (Mm), chicken (Gg) rat (Rn) and zebrafish (Dr) TMT1A. Conserved glycines in the GxGxG domain are denoted by the red asterisks. Conserved aspartic acid at position 98 is denoted by a green asterisk. (B) Amino acid sequence homology in TMT1A from five different species. (C) Phylogenic analysis of TMT1A homologs. (D) 3D predicted modeling of Homo sapiens TMT1A. (E) Comparative 3D modeling of TMT1A from Hs (green), Mm (violet), Gg (yellow), Dr (orange) and Rn (blue). Protein backbone is shown in ribbon format. S-adenosylmethionine (SAM) is modeled in black, while conserved SAM-binding residues are displayed in stick format. (F) Immunoblot analysis of HEK-293 cells transfected to express TMT1A from humans, zebrafish, chickens, mice, or rats.

### 3.2. HEK-293 cells overexpressing TMT1A homologs are selectively resistant to thiol-containing HDACis

To determine if the TMT1A homologs from other vertebrates would be able to methylate thiols, we transfected HEK-293 cells to express human TMT1A as well as mouse, rat, chicken, and zebrafish homologs and selected clones with relatively high expression of the enzyme (Fig. 1F). Predicted molecular weights for human (Hs), mouse (Mm), rat (Rn), chicken (Gg) or zebrafish (Dr) TMT1A are approximately 22-28 kDa and the bands appeared to be approximately the same size, with zebrafish having the highest molecular weight. A second band at approximately 40 kDa was observed for humans, and a similar band was found in the mouse and rat homologs but not in the chicken or zebrafish (Fig. 1F).

Analysis of protein post-translational modification data from PhosphoSite Plus [17] of human, mouse, and rat TMT1A revealed that several amino acids are predicted to have conserved modifications. In particular, the K34, K55, K105, and K219 sites were predicted to have conserved ubiquitination modifications in human and mouse. K105-ub has been identified in several high-throughput studies [18, 19], possibility indicating its importance to TMT1A function or localization. Conversely, K211-ubiquitination and S218-phosphorylation were predicted to be conserved in mice and rats, but those modifications were not observed in humans. The presence of the ubiquitination sites may contribute to the bands observed at higher molecular weight.

Having confirmed expression of the TMT1A homologs in the transfected cells, we performed three-day cytotoxicity assays with known substrates of human TMT1A. We previously reported that overexpression of TMT1A in MCF-7 cells conferred resistance to a number of thiol-containing HDACis including romidepsin, KD-5170, and NCH-51 [9]. When we compared the resistance conferred by the TMT1A homologs, we observed resistance to all the thiol-containing drugs compared to empty vector cells, although there were differences in the extent of resistance among the homologs (Fig. 2A). The highest levels of romidepsin resistance were found in cells expressing human, mouse, and rat TMT1A, while cells expressing the chicken and zebrafish homologs were less resistant. Chicken TMT1A conferred less resistance to KD-5170 (Fig. 2A) and the rat homolog conferred less resistance to NCH-51 compared to the other homologs (Fig 2A). Mouse and rat TMT1A conferred equal levels of resistance to NCH-51 compared to human. As expected, we did not observe any resistance to belinostat (Fig. 2A), a histone deacetylase inhibitor that does not have a thiol as the zinc-binding group.

**Fig. 2.**
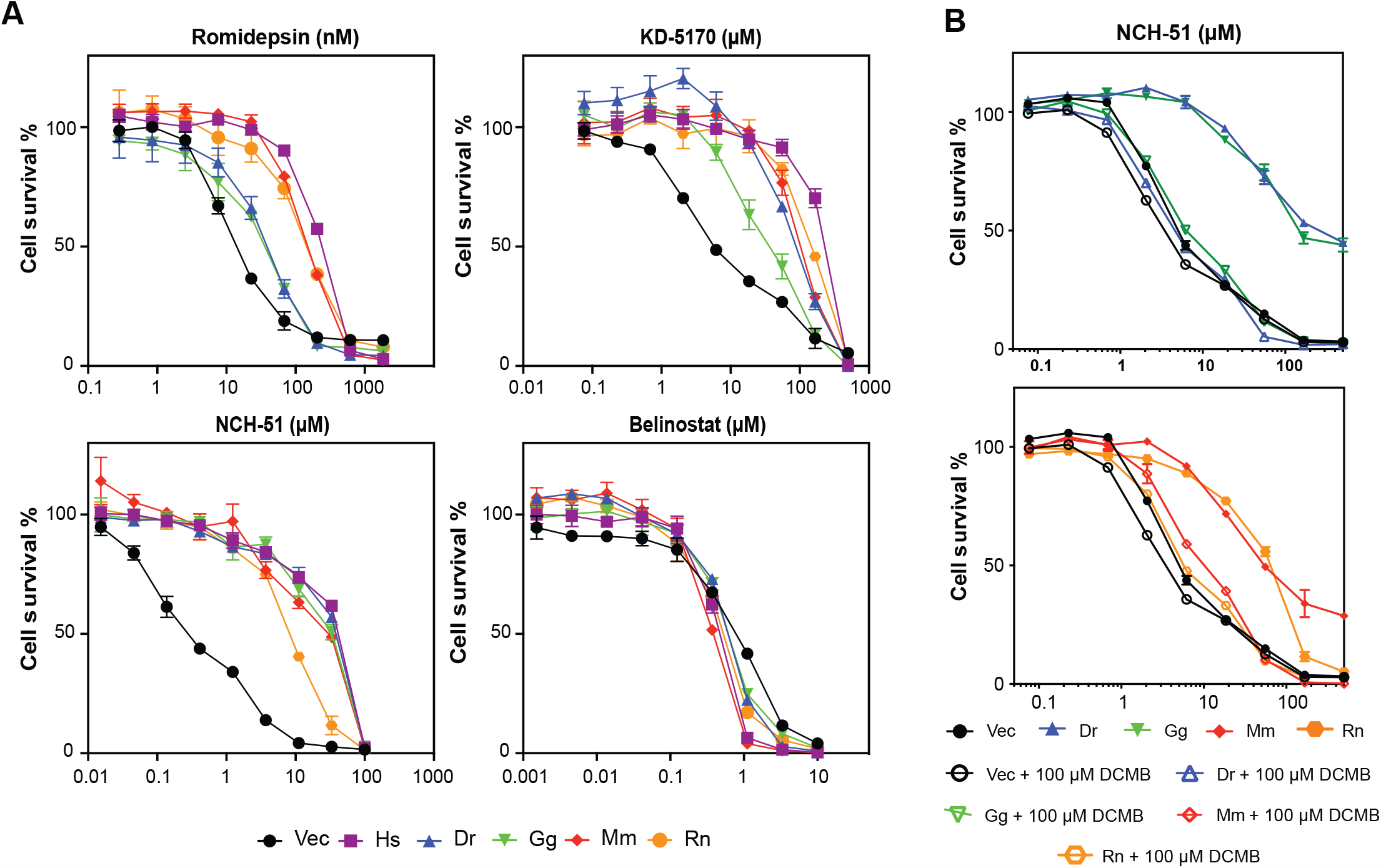
Overexpression of TMT1A homologs confers resistance to thiol-based HDACis (A) Three-day cytotoxicity assays were performed with romidepsin, KD-5170, NCH-51 or belinostat on empty vector cells (Vec) or cells expressing human (Hs), zebrafish (Dr), chicken (Gg), mouse (Mm), or rat (Rn) TMT1A. Results from one of at least three experiments are shown. (B) Three-day cytotoxicity assays were performed with NCH-51 in the presence or absence of 100 μM DCMB on cells expressing empty vector (black), or TMT1A from zebrafish (Dr), chicken (Gg), mouse (Mm), or rat (Rn).

### 3.3. DCMB reverses resistance mediated by TMT1A homologs

The phenylethanolamine-N-methyltransferase (PNMT) inhibitor DCMB was previously shown to inhibit TMT1A activity [7, 8]. To determine if DCMB could also inhibit the activity of the TMT1A homologs, cytotoxicity assays were performed with NCH-51 in the presence or absence of 100 μM DCMB, and as we previously showed, this was able to reverse TMT1A-mediated resistance in transfected cells [9]. When DCMB was added to NCH-51, resistance conferred by any of the homologs was completely reversed (Figure 2B).

### 3.4. TMT1A homologs blunt the effects of HDACi treatment

Treatment with HDACis is known to result in increased global histone acetylation and upregulation of many genes, including *CDKN1A* which codes for p21. Upregulation of p21 is known to cause cell cycle arrest and treatment with HDACis results in either a G1/G0 and/or G2/M arrest [20, 21]. To determine if the homologs of TMT1A would provide protection from the effects of HDACi treatment, cells expressing empty vector or the TMT1A homologs were treated with 10 ng/ml romidepsin (Dp), 10 μM KD-5170 (KD) or 25 μM NCH-51 (NCH) for 24 h and cell cycle analysis was performed. As shown in Figure 3A, empty vector cells (Vec, top row) demonstrate an increase in G2/M cells when treated with any of the HDACis, while cells expressing rat TMT1A (Rn, bottom row) did not display any changes in the cell cycle. Results with all the HDACis are summarized in Figure 3B and we found that treatment of cells expressing any of the TMT1A homologs with HDACis resulted in cell cycle changes that were less drastic than those observed in the empty vector cells. These results are in agreement with the cytotoxicity data, which indicated that overexpression of any of the homologs conferred resistance to thiol-based HDACis.

**Fig. 3.**
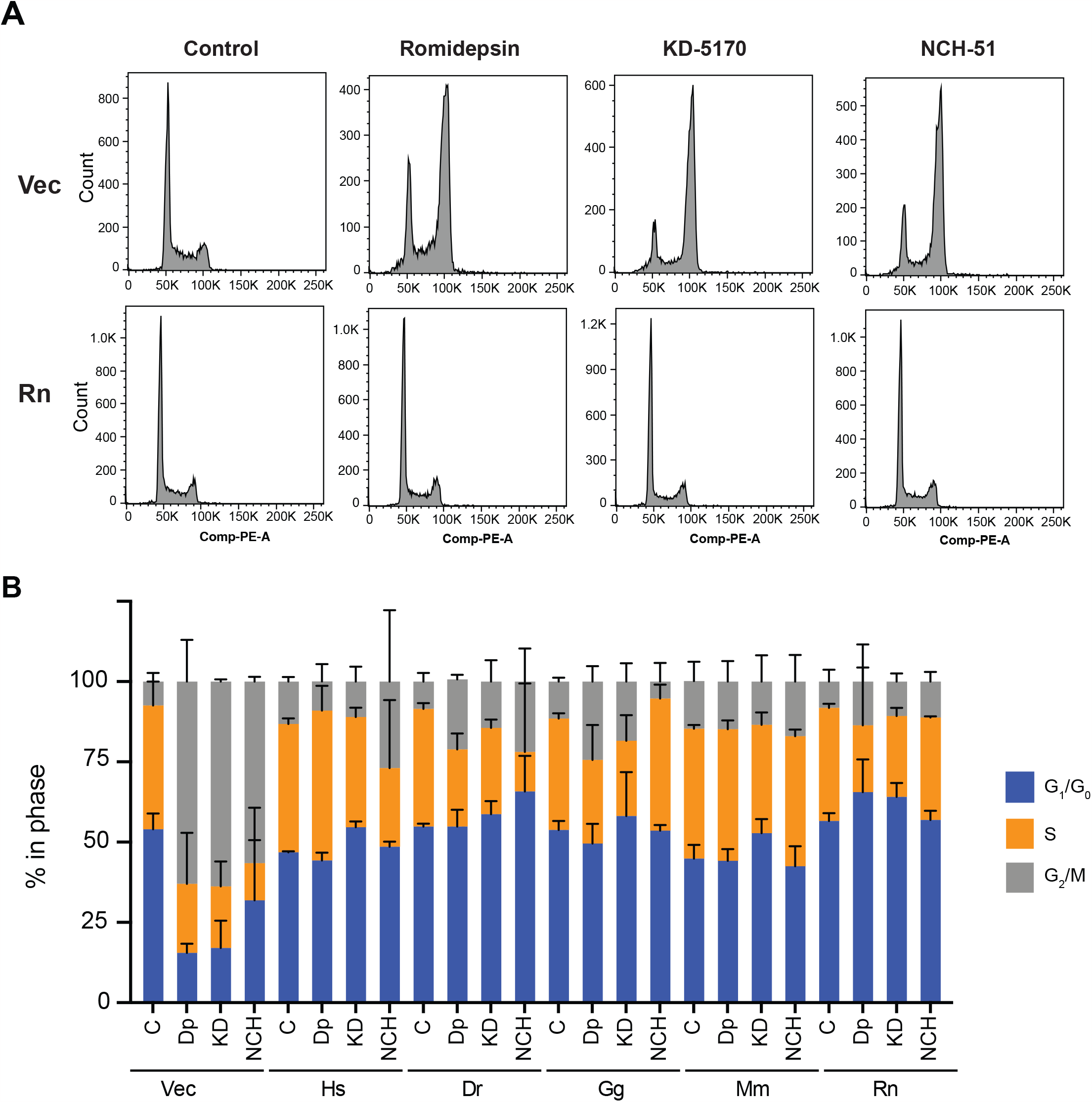
TMT1A homologs prevent cell cycle effects of HDACi treatment (A) Cell cycle analysis was performed on empty vector (Vec) cells or cells expressing rat (Rn) TMT1A after a 24 h treatment with romidepsin (10 ng/ml), KD-5170 (10 μM), or NCH-51. (B) The percentage of cells in each phase of the cell cycle was determined from empty vector cells or cells expressing any of the TMT1A homologs after the treatments outlined in (A) using ModFit LT v 5.0. Data compiled from 3 independent experiments were used to generate the graph. Standard deviation is indicated by error bars.

As treatment with HDACis is known to cause increased p21 expression as well as increased histone acetylation [22], we next examined the effect of HDACi treatment on histone acetylation and p21 expression after a 24h treatment with 10 ng/ml romidepsin. Treatment of empty vector cells with romidepsin resulted in an increase in acetylated histone H3 and p21 (Fig. 4), while this effect was blunted in cells expressing any of the TMT1A homologs. These results suggest that expression of any of the TMT1A homologs led to resistance to the effects of thiol-based HDACis most likely due to methylation of the zinc-binding thiol.

**Fig. 4.**
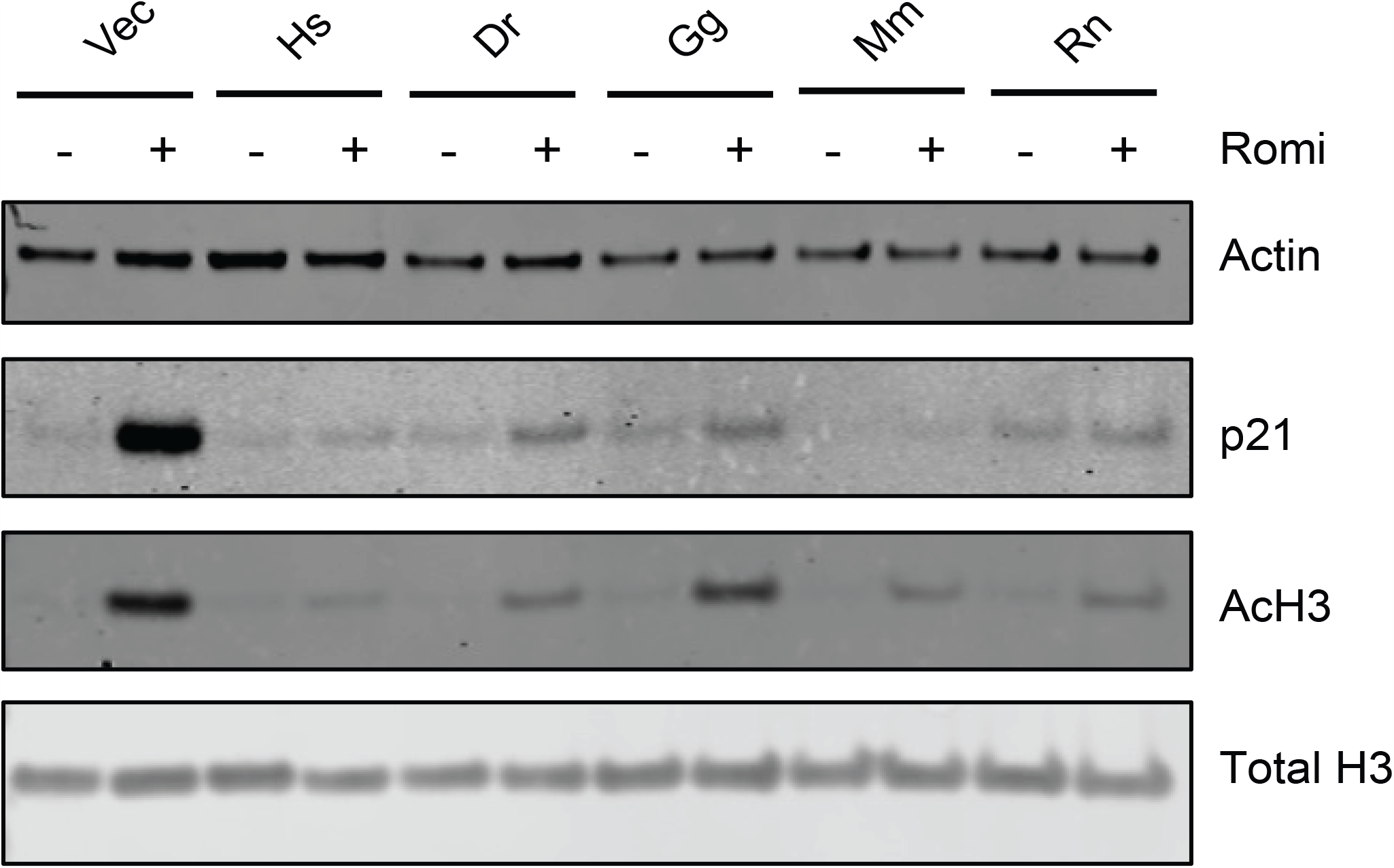
TMT1A homologs protect from downstream effects of HDACi treatment. Cells were treated with 10 ng/ml romidepsin for 24 h after which proteins were extracted, subjected to PAGE, and transferred to nitrocellulose membranes as detailed in the Materials and Methods. Blots were subsequently probed for p21, acetylated histone H3, total histone H3, and beta actin.

### 3.5. Detection of dimethylated romidepsin in culture medium of cells expressing TMT1A homologs

Romidepsin is reduced intracellularly to yield the active form of the drug that has two thiol groups, one of which coordinates with a zinc ion to inhibit HDACs [10]. We previously found that when cells that overexpress TMT1A are treated with romidepsin, dimethylated romidepsin can be detected in the culture medium [9]. To determine if TMT1A homologs from different species could also methylate romidepsin, empty vector cells and cells expressing the TMT1A homologs were incubated with 10 μM romidepsin for 24 h and both unmodified and dimethylated romidepsin were measured by LC-MS/MS. Cells expressing zebrafish or chicken TMT1A had the lowest proportion of dimethylated romidepsin relative to total romidepsin, while cells expressing the mouse or rat isoforms had levels comparable to cells expressing human TMT1A (Fig 5). These results are in accordance with cytotoxicity assay data, which show that cells expressing zebrafish or chicken TMT1A were less resistant to romidepsin.

**Fig. 5.**
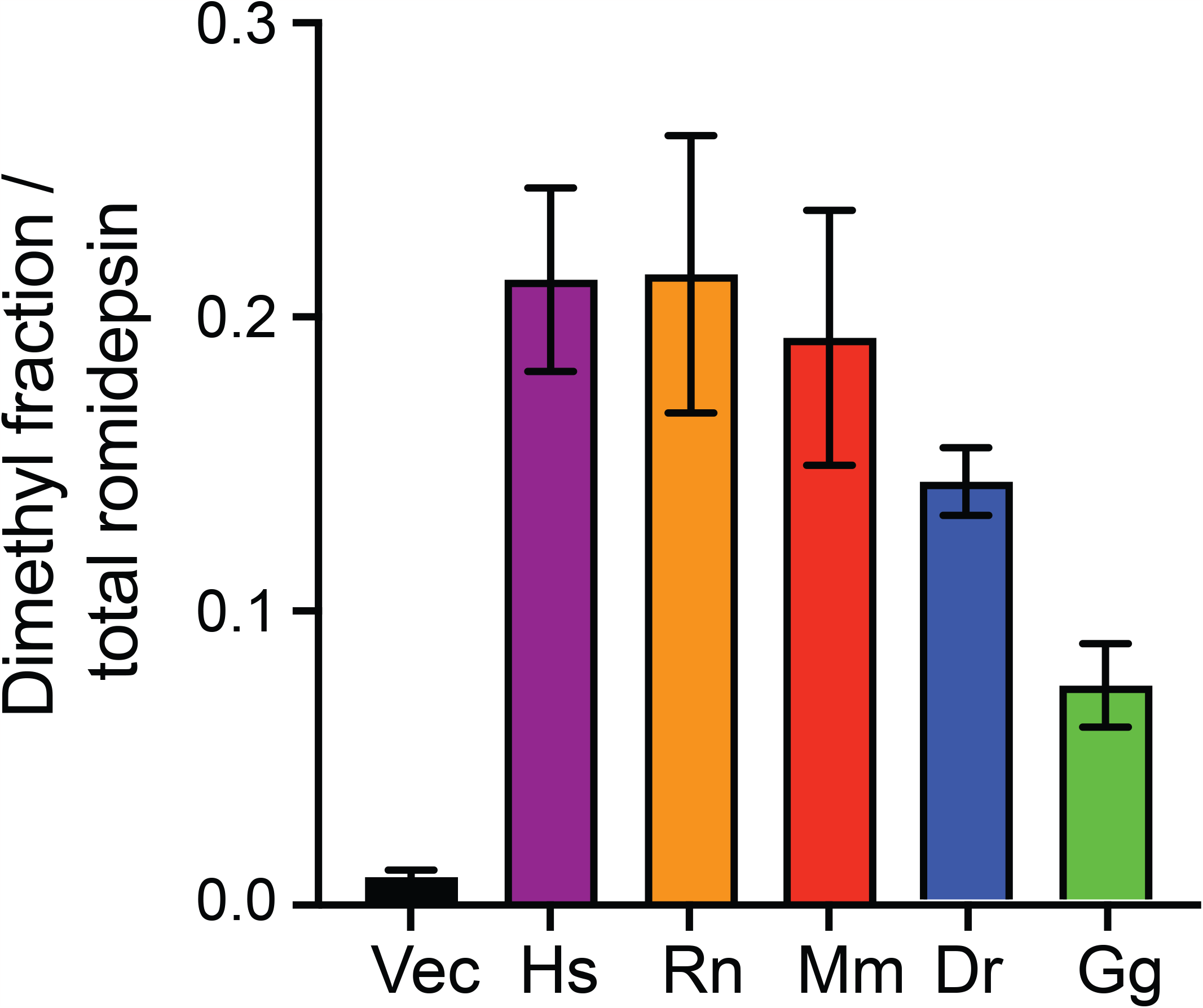
Detection of dimethylated romidepsin in culture media of transfected cells. Cells were plated and allowed to attach overnight after which 10 μM romidepsin was added to culture media and incubated for 24 h. Media was collected and mass spectrometry was performed to quantify the amount of dimethylated romidepsin relative to total romidepsin, as detailed in the Materials and Methods section.

## 4. DISCUSSION

Early studies with liver microsomes suggested the existence of a thiol methyltransferase that could methylate thiol groups on molecules [4-6]. Recently, TMT1A and TMT1B have been identified as thiol methyltransferases capable of methylating molecules such as hydrogen sulfide, captopril and thospironolactone [7, 8]. We also recently reported that TMT1A and TMT1B can methylate and inactivate HDACis with a thiol as the zinc-binding group [9]. To determine if the thiol methyltransferase activity of these enzymes is evolutionarily conserved, we transfected HEK-293 cells to express human, zebrafish, chicken, mouse, or rat TMT1A. TMT1B was not found in the chicken or zebrafish. We found that overexpression of any TMT1A isoform was able to confer resistance to any of the HDACis studied that has a thiol as the zinc-binding group. Additionally, overexpression of any of the TMT1A forms protected the cells from downstream effects of HDACi treatment, such as cell cycle arrest and increased levels of histone acetylation and p21. We thus conclude that the thiol methyltransferase activity is conserved across species and suggest that animal models may indeed aid in the determination of the physiological role of TMT1A.

Previous work on the conservation of enzyme function has shown that when the pairwise sequence identity of two proteins is above approximately 40%, they are evolutionarily related [23, 24], while sequence identity above approximately 70% can be linked to conservation of enzyme function [25]. Further, conservation of catalytic residues is expected to be highest among homologs that catalyze the same reaction, followed by enzymes that perform the same reaction but on different substrates [26]. When we compared the amino acid sequences of the TMT1A homologs, we observed high pairwise sequence identity (Figure 1B) and near universal conservation of the catalytic residues, strengthening our hypothesis that these homologs would all be able to act on thiol-containing HDAC inhibitors. Similarly, when we perform 3D structure prediction with AlphaFold, we observe that TMT1A proteins from different species all adopt the conserved seven-beta-strand (7BS) MTase fold and show good structural alignment of the conserved *S*-adenosyl methionine (SAM)-binding domain (Figure 1E). However, we note that the predicted structure may differ from the actual enzyme structure.

As TMT1A and TMT1B are poorly described, little is known about their physiological function. Interestingly, although they are thiol methyltransferases, they do not seem to methylate thiols on endogenous compounds such as glutathione or cysteine [7, 8]. However, both enzymes are able to methylate hydrogen sulfide, which has been demonstrated to have multiple functions in the cell [27] and may be dysregulated in cancer. Elevated levels of methanethiol have been found in patients diagnosed with oral squamous cell carcinoma [28]. In this study, we demonstrate that these enzymes share high sequence and structure conservation, and that the thiol methyltransferase activity of TMT1A is conserved across different species. Thus, generation of animals lacking either or both TMT1A and TMT1B could shed light on the normal physiological roles of these methyltransferases.

Although the normal physiological functions of TMT1A and TMT1B are unclear, one can hypothesize a role in drug metabolism. We found expression of TMT1A in the liver, breast epithelium, glial cells, and Leydig cells of the testis and TMT1B in the liver, intestinal enterocytes, and neurons [9], consistent with a protective role. TMT1A expression was also found in Bergmann glial cells, suggesting a role in the cerebrospinal fluid-brain barrier. Russell and colleagues also reported high levels of TMT1A and TMT1B in human liver microsome samples and protein expression of both enzymes correlated with thiospironolactone methylation activity [8]. Interestingly, methyltransferase activity in samples varied by 8-fold [8], suggesting potential interpatient variability in the metabolism of thiol-containing drugs. Although mutations of TMT1A or TMT1B could affect their activity, further work needs to be done to support this hypothesis.

## Author contributions

**J.M. González Dalmasy**: Investigation, methodology, data analysis, visualization, validation, writing–original draft, writing–review and editing. **C.M. Fitzsimmons**: Conceptualization, investigation, methodology, data analysis, visualization, writing– original draft, writing–review and editing. **A.J. Perciaccante:** Investigation, methodology, data analysis, writing–review and editing. **C.P. Jewell:** Investigation, methodology, data analysis, writing–review and editing. **W.J.E. Frye:** Investigation, methodology. **L.M. Jenkins**: Investigation, methodology, data analysis, writing–review and editing. **P.J. Batista:** Supervision, project administration, writing–review and editing. **R.W. Robey:** Conceptualization, investigation, methodology, data analysis, validation, visualization, writing–original draft, project administration, writing– review and editing. **M.M. Gottesman:** Resources, supervision, funding acquisition, project administration, writing–review and editing.

## Data availability

All data are presented in the paper and its figures.

## ACKNOWLEDGEMENTS

We appreciate the editorial assistance of George Leiman. Mention of trade names, commercial products, or organizations does not imply endorsement by the U.S. Government. This work was supported by the Intramural Research Program of the National Cancer Institute (NCI) in the National Institutes of Health. CMF was partially supported by a postdoctoral fellowship from the American Cancer Society (PF-19-157-01-RMC).

